# Translocation of chloroplast NPR1 to the nucleus in retrograde signaling for adaptive response to salt stress in tobacco

**DOI:** 10.1101/2021.03.24.436779

**Authors:** So Yeon Seo, Ky Young Park

## Abstract

Chloroplasts play a pivotal role in biotic and abiotic stress responses, accompanying changes in the cell reduction/oxidation (redox) state. Chloroplasts are an endosymbiotic organelle that sends retrograde signals to the nucleus to integrate with environmental changes. This study showed that salt stress causes the rapid accumulation of the nonexpressor of pathogenesis-related genes 1 (NPR1) protein, a redox-sensitive transcription coactivator that elicits many tolerance responses in chloroplasts and the nucleus. The transiently accumulated chloroplast NPR1 protein was translocated to the nucleus in a redox-dependent manner under salinity stress. In addition, immunoblotting and fluorescence image analysis showed that chloroplast-targeted NPR1-GFP fused with cTP (chloroplast transit peptide from RbcS) was localized in the nucleus during the responses to salt stress. Chloroplast functionality was essential for retrograde translocation, in which the stomules and cytoplasmic vesicles participated. Treatments with H_2_O_2_ and an ethylene precursor enhanced this retrograde translocation. Compared to each wild-type plant, retrograde signaling-related gene expression was severely impaired in the *npr1-1* mutant in Arabidopsis, but enhanced transiently in the *NPR1-Ox* transgenic tobacco line. Therefore, NPR1 might be a retrograde signaling hub that improves a plant’s adaptability to changing environments.

## Introduction

How organelles communicate with the nucleus to coordinate genetic programs and cellular functions is a fundamental question of plant physiology and cell biology (Pfannschmidt et al., 2020). Reduction/oxidation (redox)-associated signaling is an essential component of responses to environmental stresses and pathogen attack in all organisms, including plants (Munné-Bosch et al., 2013), in which stress-related reactive oxygen species (ROS) and redox information are principally accumulated in chloroplasts. Disturbance to photosynthetic activity in chloroplasts under environmental stresses or pathogen attack creates an oxidative environment, facilitating signaling by oxidation of protein cysteine residues (van der Reest et al., 2018). Therefore, ROS-sensitive proteins function as redox switches in response to abiotic/biotic stress. It was recently reported that a ROS-mediated redox cascade is involved in communication between chloroplasts and the nucleus through retrograde signaling, resulting in protective mechanisms and modulation of hormone biosynthesis (Chan et al., 2016; Müllineaux et al., 2020). Although retrograde signals from chloroplasts to the nucleus for adaptive responses during chloroplast development and under environmental stresses have been explored recently, the translocation mechanism of metabolites or proteins is still poorly understood. Proteins with redox-sensitive cysteine residues may function as redox sensors and retrograde signaling switches through redox-sensitive post-translational modifications (PTM) (Smirnoff and Arnaud, 2019; Mata-Pérez and Spoel, 2019).

The nonexpressor of pathogenesis-related genes 1 (NPR1) protein is a transcription coactivator and a master regulator of plant immunity with salicylic acid (SA)-mediated defense responses and systemic acquired resistance (SAR) in Arabidopsis (Mou et al., 2003). NPR1 proteins sense cytoplasmic changes in SA-dependent redox status during innate immune responses (Tada et al., 2008). Pathogen-induced SA triggers alteration of the cellular reduction potential, thereby reducing the cytoplasmic NPR1 tetramer into a monomer via breakage of disulfide bonds, after which the NPR1 monomer is imported into the nucleus to function as a coactivator of gene transcription in SAR (Spoel et al., 2009). On the basis of our previous results, although NPR1 is a transcriptional coactivator, a large amount of NPR1 is present in the chloroplasts under salt stress (Seo et al., 2020).

## Results

### Translocation of NPR1 from chloroplasts to the nucleus under salt stress

In a previous study, we generated stable transgenic tobacco plants expressing a fusion construct of full-length *NPR1* combined with the green fluorescent protein (*GFP*) driven by a 0.8-kb region of the *NPR1* promoter (*pNPR1::NPR1-GFP*) or the *CaMV 35S* nuclear promoter (*p35S::NPR1-GFP*) (Seo et al., 2020). With increase in salt concentration, translocation of NPR1 to the chloroplasts under salt stress increased significantly (Supplemental Figure 1A and 1B). In the present study, we further investigated the cellular partitioning of NPR1 under salt stress using a confocal laser scanning microscope (STELLARIS 8; LEICA, Germany). Peak NPR1-GFP fluorescence in chloroplasts was observed at 12 h, then declined rapidly to similar to the initial intensity at 24 h, whereas fluorescence was strongly observed in the nucleus at 24 h, in tobacco mesophyll protoplasts of *pNPR1::NPR1-GFP* transformants under salt stress (Figure 1A and Supplemental Movie1). We noted that a large amount of NPR1 was present in the nucleus at the time that NPR1 disappeared from the chloroplast after 24 h of salt stress, hence additional experiments were performed to further explore this relationship. In particular, we investigated whether NPR1 in the nucleus is imported from the cytoplasm or whether chloroplast NPR1 is translocated to the nucleus. Notably, NPR1-GFP vesicles of various sizes were observed in the cytoplasm at 6 h under salt stress (Figure 1B). Cytoplasmic NPR1 condensates are observed in Arabidopsis leaves after treatment with SA, which performs an essential function in mediating protein homeostasis and cell survival during the plant immune response (Zavaliev et al., 2020). Therefore, we investigated the function of cytoplasmic NPR1 vesicles in salt-stressed tobacco leaves.

**Figure 1.**
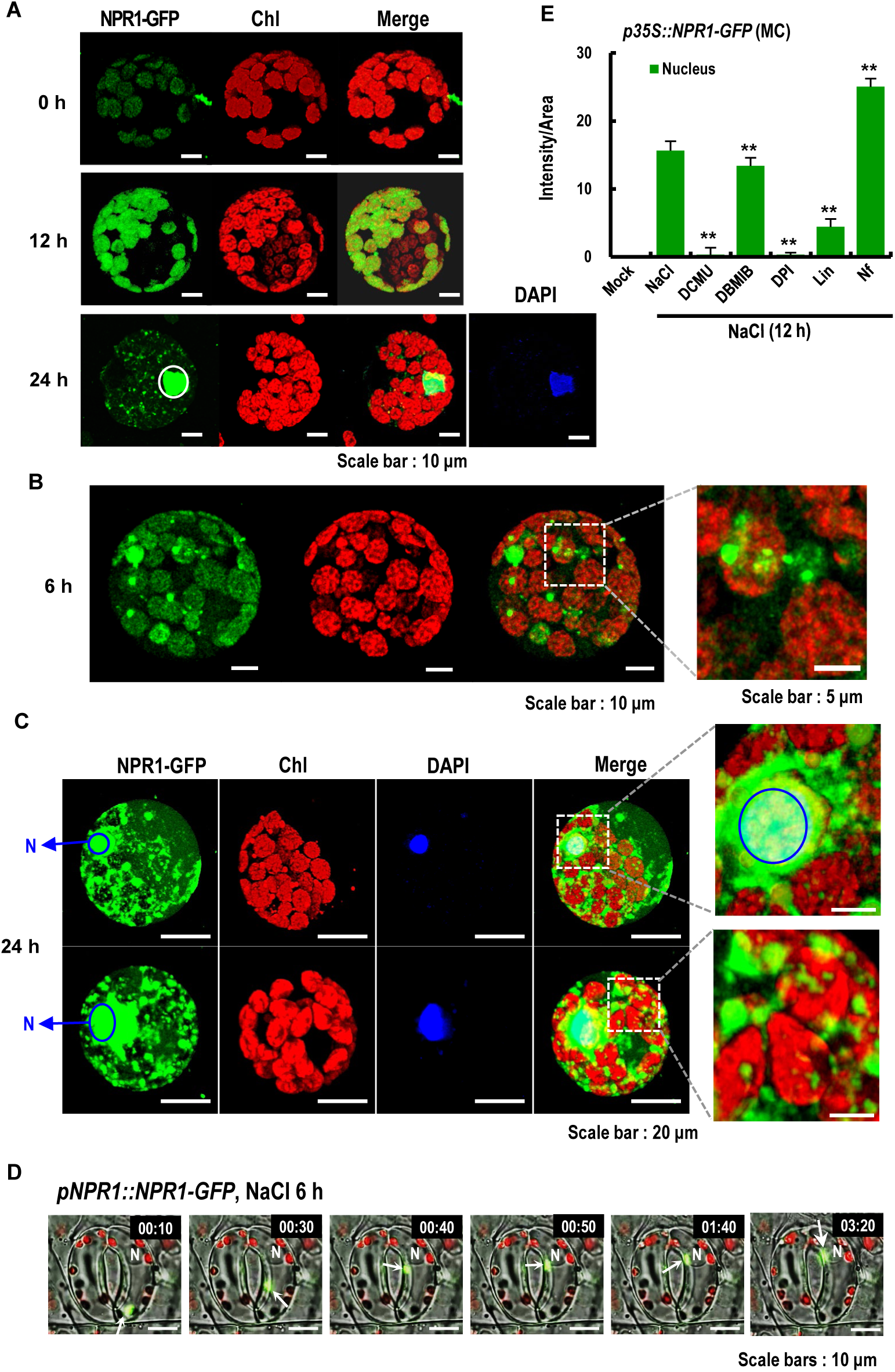
Intracellular localization of GFP-tagged NPR1 in leaf cells under salt stress. **(A)** Confocal laser scanning microscopy (CLSM) images of GFP fluorescence in mesophyll protoplasts from 6-week-old *pNPR1::NPR1-GFP* transgenic plants after salt stress with 200 mM NaCl. Images of GFP fluorescence (green) and chlorophyll autofluorescence from chloroplasts (red) are merged in the third column. DAPI staining (blue) was in the fourth column of the last row. **(B)** Cytoplasmic vesicles containing NPR1-GFP observed in the cytoplasmic region. NPR1-GFP is contained in green vesicles of various sizes. Cytoplasmic vesicles within the area enclosed by the dotted line are highlighted. **(C)** Localization of NPR1-GFP in the nucleus at 24 h after salt stress treatment. Highlights of the nuclear NPR1-GFP, which is merged with autofluorescence and DAPI images (upper right). In the enlarged image, the nucleus area within the dotted line is highlighted. **(D)** Snapshots of rapidly moving cytoplasmic vesicles containing NPR1-GFP in guard cells from the abaxial epidermis of *pNPR1::NPR1-GFP* transgenic plants at 6 h after salt stress treatment. White arrows indicate vesicles. N: nucleus. **(E)** Fluorescence intensity of NPR1-GFP in the nucleus from *p35S::NPR1-GFP* transgenic plants under salt stress. Inhibitors were co-treated with salt stress. Inhibitors: 3-(3,4-dichlorophenyl)-1,1-dimethylurea (DCMU), 2,5-dibromo-3-methyl-6-isopropylbenzoquinone (DBMIB), diphenyleneiodonium (DPI), lincomycin (Lin), norflurazone (Nf). An asterisk indicates a significant difference between transformants treated with salt only and transformants co-treated with salt and other chemicals (\*\**P* < 0.01).

Although Arabidopsis NPR1 is predominantly sequestered in the cytoplasm as a high-molecular-weight oligomeric complex, upon pathogen infection the cellular reduction potential is changed by cytoplasmic SA and thioredoxin, which results in partial reduction of the NPR1 oligomer to a monomer and its translocation to the nucleus where it functions as a coactivator of TGA transcription factors (Després et al., 2003). However, tobacco NPR1 was rapidly imported into chloroplasts after pathogen infection with *Phytophthora parasitica* var. *nicotianae*, which was followed by NPR1 translocation to the nucleus (Supplemental Figure 1C and 1D). These results implied that tobacco NPR1 undergoes an intermediate process in which NPR1 is transiently sequestered in chloroplasts during the resistance response to virulent pathogens. Chloroplasts are organelles that produce large amounts of ROS in the early stages of stresses, thus redox-sensitive chloroplast proteins are ideal candidates as redox sensors or signaling molecules. In this proposed mechanism, chloroplasts determine the current environmental state and produce diverse signals informing the nucleus about the functionality of the photosynthetic apparatus, which is defined as retrograde signaling (Pfalz et al., 2012).

In the response to salt stress, tobacco NPR1 was gradually localized in the chloroplasts, and at the same time, many vesicles gradually developed around the chloroplasts at 6 h (Figure 1B). After 24 h of salt stress, NPR1 was localized predominantly in and around the nucleus and the cytoplasm (Figure 1C). When the blue nuclear fluorescence from DAPI staining was merged with the green-fluorescent NPR1 signal, the blue-green signal was clearly apparent, suggesting that NPR1 was localized in the nucleus (Figure 1C, upper panel). In the merged images, the green band surrounding the sky-blue region indicated that NPR1 was localized in the perinuclear region. Interestingly, the observations implied that a large amount of NPR1-GFP was clustered around the nucleus. The rapid movement of vesicle-like structures containing NPR1 was also observed in mesophyll protoplasts under salt stress (Supplemental Movie 2). Considering the temporal changes in subcellular localization, perinuclear aggregation of NPR1-GFP suggests that NPR1 likely exits the chloroplasts and approaches the nucleus during the stress response. Taken together, these observations implied that NPR1 may be involved in chloroplast-to-nucleus retrograde signaling.

To further investigate whether NPR1 moves from chloroplasts to the nucleus, the *pNPR1::NPR1-GFP* transformant was subjected to salt stress and movement of NPR1-GFP protein in guard cells of the leaf was explored using a fluorescence microscope (Thunder Imager, LEICA, Germany). Surprisingly, NPR1 vesicles approaching the nucleus proceeded to fuse with the nucleus, and then NPR1-GFP fluorescence rapidly disappeared, which indicated that the NPR1 protein was imported into the nucleus (Figure 1D and Supplemental Movie 3). Although stress-induced retrograde signaling pathways involving chlorophyll intermediates, ROS, metabolites, and transcription factors in chloroplast-to-nucleus communication have recently been identified, the molecular machineries of such retrograde signaling pathways are incompletely understood (Chan et al., 2016). The present study presents the first observation that proteins are released from chloroplasts in vesicle-like structures and directly access the nucleus during the stress response, although the direct transfer of H_2_O_2_ and SA through stromules or chloroplast–nucleus complexes has been observed previously (Exposito-Rodriguez et al., 2017; Caplan, 2015).

Next, we examined whether NPR1 translocation to the nucleus is affected by altering chloroplast conditions, in which the chloroplast NPR1 abundance is dependent on its oxidative status (Seo et al., 2020). We previously reported that plant cells under salt stress rapidly up-regulated the redox-sensitive NPR1 protein, which is imported to chloroplasts for induction of protective responses as chaperones and antioxidants with lower chloroplastic ROS accumulation.

The compounds 3-(3,4-dichlorophenyl)-1,1-dimethylurea (DCMU) and 2,5-dibromo-3-methyl-6-isopropylbenzoquinone (DBMIB) inhibit the photosynthetic electron transport chain in photosystem II (PSII) (Mühlenbock et al., 2008). DCMU increases the pool of oxidized plastoquinone (PQ), whereas DBMIB increases the pool of reduced PQ. Under co-treatment with salt stress, DCMU almost completely inhibited NPR1 accumulation in the nucleus, but DBMIB caused weak inhibition, in comparison with stress alone (Figure 1E). Based on the finding that DCMU completely blocked ROS accumulation in the chloroplast stroma (Exposito-Rodriguez et al., 2017), it was concluded that DCMU completely prevented NPR1 import to the nucleus because ROS were not produced in the chloroplasts.

Next, diphenyleneiodonium (DPI), an inhibitor of NADPH oxidase activity (Seo et al., 2020), was administered to leaves of *p35S::NPR1-GFP* transgenic plants under salt stress. The DPI-dependent inhibition of NADPH oxidase activity resulted in reduction of its products, including ROS, in chloroplasts and other cellular compartments. As expected, DPI treatment completely prevented NPR1 accumulation in the nucleus (Figure 1E). Lincomycin (LIN), a translation inhibitor in chloroplasts (Kim and Mullet, 2003), significantly reduced the amount of nuclear NPR1 (Figure 1E). Norflurazone, a compound inducing strong photo-oxidation and subsequent plastid dysfunction (Park et al., 2017), caused prominent nuclear NPR1 accumulation in leaf protoplasts (Figure 1E). This result suggested that norflurazone-induced photo-oxidation enhanced the translocation of NPR1 to the nucleus. Taken together, the changes in oxidative status and protein translation in chloroplasts might impact on the nuclear accumulation of NPR1 under stress.

### Changes in NPR1 translocation to the nucleus according to chloroplast condition

Next, we examined whether NPR1 targeted to chloroplasts migrates to the nucleus and affects stress tolerance. Only a small portion of the total chloroplast proteome, which lacks chloroplast transit peptides (cTP), is nucleus-encoded and synthesized on cytosolic ribosomes, and thus enters internal chloroplast compartments (Armbruster et al., 2009). Some non-cTP chloroplast proteins may be localized to the stroma through the endoplasmic reticulum (ER)-dependent chloroplast targeting pathway (Nanjo et al., 2006). Although tobacco NPR1 does not possess cTP and signal peptides, NPR1-GFP fluorescence was detected in chloroplast stroma, as evidenced by the yellow color in the merged image with chlorophyll autofluorescence (Figure 1A). To monitor the movement of chloroplast-targeted NPR1, we attached the *cTP* sequence (79 amino acid residues) of tobacco *RbcS* (GenBank accession AY220079) to the 5′ end of *NPR1-GFP* to generate *p35S::cTP-NPR1-GFP* transgenic tobacco.

Surprisingly, chloroplast-targeted NPR1-GFP with cTP was observed in the nucleus of mesophyll cells and guard cells after salt stress (Figure 2A and 2B), implying that chloroplast NPR1 moved to the nucleus under salt stress, which can be referred to as retrograde communication. Relatively constant amounts of cTP-NPR1-GFP protein accumulated in chloroplasts of *p35S*-driven *cTP-NPR1-GFP* transgenic plants under salt stress, indicating that cTP-NPR1-GFP was constitutively imported into chloroplasts mediated by the transit peptide (Supplemental Figure 2A). Although NPR1 abundance was constitutively maintained in the chloroplasts, the cTP-attached NPR1 abundance was lower in the nucleus and peaked at 9 h of salt stress. These results implied that chloroplastic NPR1 was realistically translocated to the nucleus during the response to salt stress. LIN almost entirely blocked accumulation of nuclear NPR1, which is expected to originate from chloroplasts, in cTP-fused NPR1-GFP transgenic plants (Supplemental Figure 2B).

**Figure 2.**
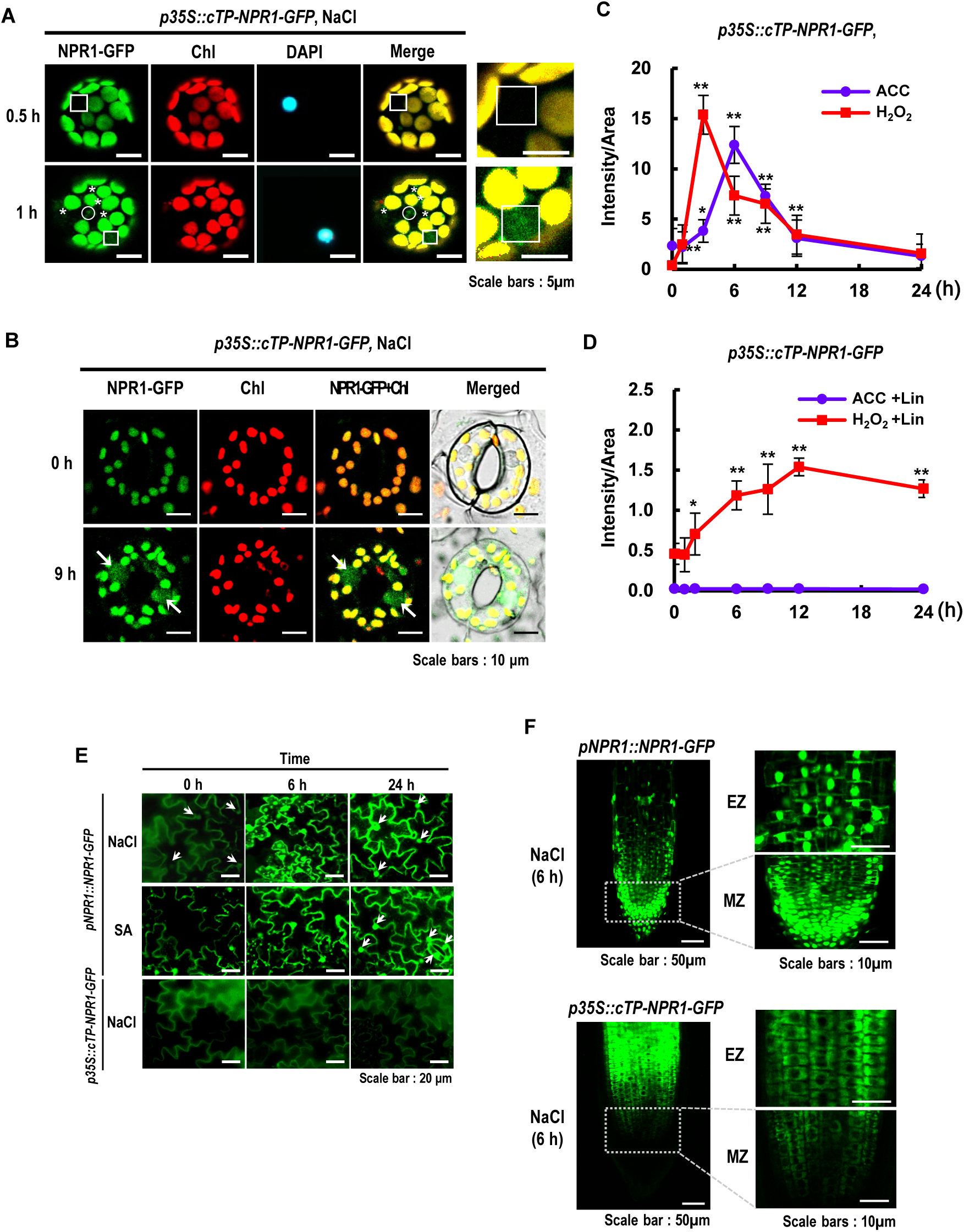
Nuclear import of NPR1-GFP under salt stress. **(A)** CLSM images of NPR1-GFP fluorescence in salt-stressed protoplasts of *p35S::cTP-NPR1-GFP* transgenic plants. Enlarged CSLM images in the last column. White box indicates the whole nucleus. White circle and asterisk indicate cytoplasmic vesicles and chloroplast protrusions, respectively. **(B)** CLSM images of NPR1-GFP in salt-stressed guard cells of *p35S::cTP-NPR1-GFP* transgenic plants. Arrows indicates the whole nucleus. **(C and D)** Fluorescence intensity of GFP in the nucleus of mesophyll protoplasts from *p35S::cTP-NPR1-GFP* transgenic plants. GFP fluorescence was photographed in mesophyll protoplasts of transgenic plants after application of H_2_O_2_ or ACC, and then GFP intensity was quantified in the nucleus (**C**). After co-treatment of an inhibitor of Lincomycin (Lin) with H_2_O_2_ or ACC, the GFP intensity was quantified in the nucleus of protoplasts isolated from transgenic plants (**D**). An asterisk indicates a significant difference from 0 h (\**P* < 0.05, \*\**P* < 0.01). **(E)** Localization of NPR1-GFP in pavement cells of the abaxial epidermis of leaves in *pNPR1::NPR1-GFP* after salt stress (upper) and SA treatment (middle) and *p35S::cTP-NPR1-GFP* after salt stress (lower). Arrows indicate NPR1 condensates. **(F)** Localization of NPR1-GFP in intracellular compartments of roots in in *pNPR1::NPR1-GFP* (upper) and *p35S::cTP-NPR1-GFP* (lower) transgenic plants under salt stress. EZ: elongation zone, MZ: meristematic zone.

Furthermore, when exogenous H_2_O_2_ or 1-amino-1-cyclopropane carboxylic acid (ACC), a precursor of ethylene, was applied to transgenic plants (*p35S::cTP-NPR1-GFP*), nuclear NPR1 increased significantly in mesophyll protoplasts (Figure 2C). In our previous study, H_2_O_2_ and ACC application resulted in increased accumulation of chloroplast NPR1 in tobacco leaves (Seo et al., 2020). These results reinforced the conclusion that NPR1 translocation from chloroplasts to the nucleus is dependent on the ROS concentration in plants. In particular, LIN almost completely blocked NPR1 accumulation in the nucleus, despite treatment with H_2_O_2_ or ACC with salt stress (Figure 2D). These results further confirmed the involvement of chloroplasts in nuclear NPR1 abundance under salt stress. Taken together, it is implied that chloroplast-to-nucleus translocation of NPR1 is required for chloroplast functionality with protein translation.

To further investigate the role of chloroplasts in NPR1 translocation to the nucleus, we determined NPR1 subcellular localization in leaf epidermal pavement cells and root of tobacco, where small chloroplasts or leucoplasts were present, respectively (Brunkard et al., 2015). Native *pNPR1*-driven NPR1-GFP was significantly accumulated in the plasma membrane, cytoplasm, and nucleus of the pavement cells in the leaf abaxial epidermis of tobacco transgenic plants after salt stress for 6 h (Figure 2E). Although weak NPR1-GFP fluorescence was detected in untreated transgenic plants, relatively high-intensity fluorescence was observed after stress treatment, which was suggested to be newly induced by salt stress. In particular, NPR1-GFP formed cytoplasmic vesicles of various sizes, which were dispersed in the cytoplasm at 6 h of salt stress, but had almost disappeared at 24 h.

Our observation of minute cytoplasmic bodies in the tobacco leaf epidermal pavement cells after 6 h of 1 mM SA treatment (Figure 2E) was consistent with a previous report that NPR1-GFP bodies were detected in the cytoplasm and nucleus in Arabidopsis after treatment with 5 mM SA for 2 h using a transient expression assay (Zavaliev et al., 2020). The NPR1 bodies were designated SA-induced NPR1 condensates, which are enriched with defense-and stress-associated proteins and ubiquitination components, such as CUL3. The NPR1-CUL3 condensates are suggested to perform functions in SA-induced regulation of protein homeostasis. In our previous report, it was revealed that cytoplasmic NPR1 shows chaperone activity under salt stress (Seo et al., 2020). Therefore, it is considered that formation of NPR1 condensates is correlated with the chaperone activity of cytoplasmic NPR1. In the present study, NPR1 bodies of various sizes were observed in the cytoplasm at 6 h of salt stress and SA treatment, after which they were almost undetectable, and NPR1 was predominantly observed in the nucleus and plasma membrane at 24 h of salt stress and SA treatment. The minute bodies were considered to be NPR1 condensates, whereas some of the larger bodies were considered to be cytoplasmic vesicles. However, the low abundance of cTP-fused NPR1 was not changed visibly in leaf epidermal pavement cells in response to salt stress (Figure 2E, lower panel). Taken together, these results indicated that NPR1 condensate abundance differed according to the status of chloroplasts and may reinforce the contention that NPR1 is imported into chloroplasts during the stress response.

Although cTP-attached NPR1-GFP (cTP-NPR1-GFP) was strongly accumulated only in the cytoplasm of cells in the root elongation zone (EZ) of transgenic plants constitutively expressing *cTP-NPR1-GFP* under salt stress, *pNPR1*-driven NPR1-GFP without cTP was significantly accumulated in the nucleus in cells of the meristematic zone (MZ) and EZ of roots (Figure 2F, Supplemental Figure 2C and 2D). These results implied that NPR1-GFP was translocated from the cytoplasm to the nucleus in roots during the stress response regardless of chloroplast functionality, but cTP-fused NPR1 (cTP-NPR1-GFP) was not translocated to the nucleus and instead remained in the cytoplasm. Moreover, *p35S*-driven cTP-NPR1 was barely detected in the MZ of the root, suggesting that nonfunctional NPR1 with the transit peptide in the cytoplasm of MZ root cells may be completely degraded (Figure 2F). It was previously reported that nuclear-encoded RbcS with cTP was not imported into root plastids (Yan et al., 2006). Therefore, it can be assumed that cTP-NPR1, which could not migrate to the nucleus in root MZ or leaf epidermal pavement cells, is continuously degraded by proteolytic cleavage. Taken together, these results indicated that cTP presence interrupted the nuclear localization of NPR1, suggesting that the involvement of PTM in chloroplasts is required for chloroplast-to-nucleus signaling during the stress response.

Next, a transient assay was performed on tobacco protoplasts with a native *pNPR1*-driven construct in which *GFP* was fused to the N-terminus of *NPR1*. Although GFP was present at the N-terminus of NPR1, GFP-NPR1 fluorescence was strongly detected in the chloroplasts under salt stress, which showed the same pattern as NPR1-GFP (Supplemental Figure 3). However, GFP-NPR1 was not detected in the nucleus. Given GFP-NPR1 translocation only into the chloroplasts, this result indicated that NPR1 was imported into the chloroplasts independently of the N-terminus of NPR1.

Maximal photochemical efficiency and trypan blue staining for cell death indicated that *p35S*-driven overexpression of *NPR1-GFP* resulted in greater tolerance to salt stress. Compared with the wild-type (WT), transgenic plants expressing *cTP-NPR1-GFP* or *NPR1-GFP* under salt stress showed enhanced maximal photochemical efficiency of PSII (*F*_v_*/F*_m_) (Maruta et al., 2012) based on chlorophyll fluorescence measured using a PAM 2000 Photosynthesis Yield Analyzer (Walz, Germany) (Figure 3A).

**Figure 3.**
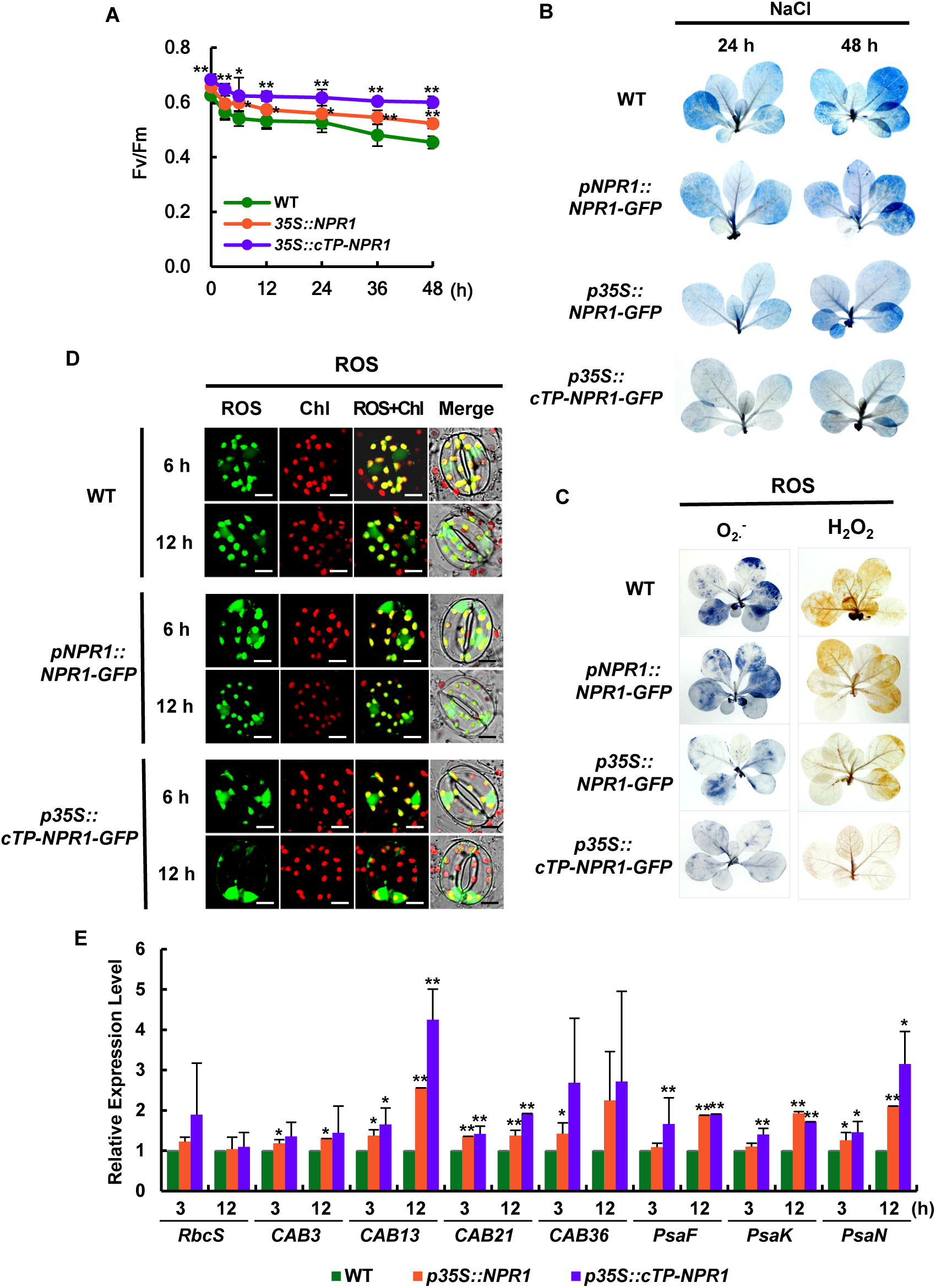
Enhancement of chloroplast-targeted NPR1 in stress resistance. **(A)** The maximal photochemical efficiency of photosystem II (*F*_v_/*F*_m_) was measured in WT, *p35S::NPR1-GFP*, and *p35S::cTP-NPR1-GFP* tobacco plants after salt stress. **(B)** Necrotic areas in salt-stressed plants were stained with trypan blue. **(C)** Histochemical analysis of ROS accumulation. Superoxide anion was detected by NBT staining (left), and H_2_O_2_ was detected by DAB staining (right). **(D)** Accumulation of ROS in cellular compartments of guard cells in WT and transgenic plants under salt stress. ROS was determined using CLSM after staining with 50 μM DCFH-DA. **(E)** Kinetics of nuclear-encoded gene transcription in WT and two transgenic plants upon salt stress. Nucleus-encoded genes: *Rbc S*, RubisCO Small subunit; *CAB3*, Chlorophyll *a*/*b*-binding protein 3; *CAB12*, Chlorophyll *a*/*b*-binding protein 12, *CAB21*, Chlorophyll *a*/*b*-binding protein 21; *CAB36*, Chlorophyll *a*/*b*-binding protein 36; *PsaF*, Photosystem I reaction center subunit III; *PsaK*, Photosystem I subunit X; *PsaN*, Photosystem I reaction center subunit XII. The relative mRNA expression levels are expressed as the mean ± SD. An asterisk indicates a significant difference between WT and transgenic plants at an indicated time (\**P* < 0.05, \*\**P* < 0.01).

Constitutive expression of *cTP-NPR1-GFP* more significantly reduced cell damage under salt stress compared with NPR1 without cTP (Fig 3B). However, in transgenic plants in which NPR1 was induced by native *pNPR1*, cell damage caused by salt stress was not notably different from that of WT plants. These results are associated with the functions of chloroplast-localized NPR1 as an antioxidant and a chaperone (Seo et al., 2020). cTP-NPR1 driven by *p35S* may have been present in the chloroplast before stress treatment and was then moved to the nucleus after processing to remove cTP, and the action of NPR1 as a transcription coactivator may have caused expression of resistance-related proteins and antioxidant enzymes. Therefore, experiments focused on the mechanism by which NPR1 enhanced resistance to salt stress were performed.

We visualized *in vivo* ROS generation with 2′,7′-dichlorofluorescin diacetate, which is a fluorogenic dye for cellular ROS including H_2_O_2_, and staining with nitro blue tetrazolium and diaminobenzidine for microscopic detection of O_2_^.−^ and H_2_O_2_, respectively. In cells with increased expression of NPR1, ROS accumulation in the chloroplasts visibly decreased compared with that of the WT under salt stress (Figure 3C). In particular, the cTP-fused NPR1 further suppressed ROS accumulation, including O_2_^.−^ and H_2_O_2_, in whole leaves of transgenic plants under salt stress (Figure 3C). However, ROS accumulation increased visibly in the nucleus from the onset of salinity stress, peaked at 6 h, and thereafter decreased in WT and *pNPR1::NPR1-GFP* plants, but a high amount of ROS was maintained in the nucleus of guard cells in *p35S::cTP-NPR1-GFP* plants (Figure 3D). These results suggested that elevated content of NPR1 in chloroplasts more efficiently caused ROS/redox regulation of chloroplast–nucleus communication in *p35S::cTP-NPR1-GFP* transformants. Elevated ROS up-regulates the expression of transcription factors and stress-related proteins in the nucleus, possibly enhancing stress tolerance.

The pattern for increase in the *F*_v_*/F*_m_ ratio was consistent with the gene expression characteristics of chloroplastic components for photosynthesis in *p35S::cTP-NPR1-GFP* transgenic plants (Figure 3E). Hydrogen peroxide-triggered retrograde signaling from chloroplasts to the nucleus plays a specific role in response to abiotic stress (Hanson and Hines, 2018) and innate immunity (Caplan, 2015). The present results also suggested that chloroplast NPR1 is a prerequisite for nuclear NPR1, which might be dependent on the oxidative status of chloroplasts. To elucidate the physiological functions of nuclear NPR1 in response to salt stress, we compared the transcription patterns of nuclear-encoded genes for photosynthesis-related proteins among WT and transgenic plants (*p35S::NPR1-GFP* and *p35S::cTP-NPR1-GFP*) upon salt stress. Real-time quantitative RT-PCR (qRT-PCR) was performed on the genes encoding RubisCO and core complex and antenna proteins of PS I and II. The transcript levels in transgenic plants compared with those in the WT were higher in almost all genes at 3 h and 12 h after salt stress (Figure 3E). In particular, transcript ratios for *p35S::cTP-NPR1-RFP* to WT increased significantly in all tested nuclear-encoded genes after 12 h of salt stress, when the NPR1-GFP protein was prominently localized in the nucleus (Figure 2A and 2B).

NPR1 is controlled by the nuclear localization sequence (NLS) at the C-terminus and functions as a transcription coactivator in the nucleus (Després et al., 2003). The NLS region at the C-terminus of NPR1 was deleted, after which a mutated construct, *p35S::NPR1(Δnls)-GFP*, was transiently co-expressed with *p35S::NPR1-cyan fluorescent protein* (*CFP*) in mesophyll protoplasts isolated from WT tobacco plants. To investigate NPR1 translocation from chloroplasts to the nucleus after salt stress, two *p35S*-driven variants [NPR1-CFP and NPR1(Δnls)-GFP] were used. The GFP or CFP fluorescence intensity was determined under salt stress after transient co-expression in leaf protoplasts (Figure 4A). The intensity of NPR1-CFP fluorescence in the chloroplasts peaked at 6 h and thereafter gradually decreased. In particular, vesicles containing only NPR1-CFP were observed around the chloroplasts, which resembled chloroplast protrusions (Figure 4B). However, NPR1(Δnls)-GFP fluorescence intensity continuously increased in chloroplasts until 36 h (Figure 4C). Chloroplast NPR1-GFP from this construct without the NLS was maintained at a much higher level compared with that of the *NPR1-CFP* construct with the NLS (Supplemental Figure 4). These results suggested that stress-induced export of NPR1 from chloroplasts was dependent with the NLS.

**Figure 4.**
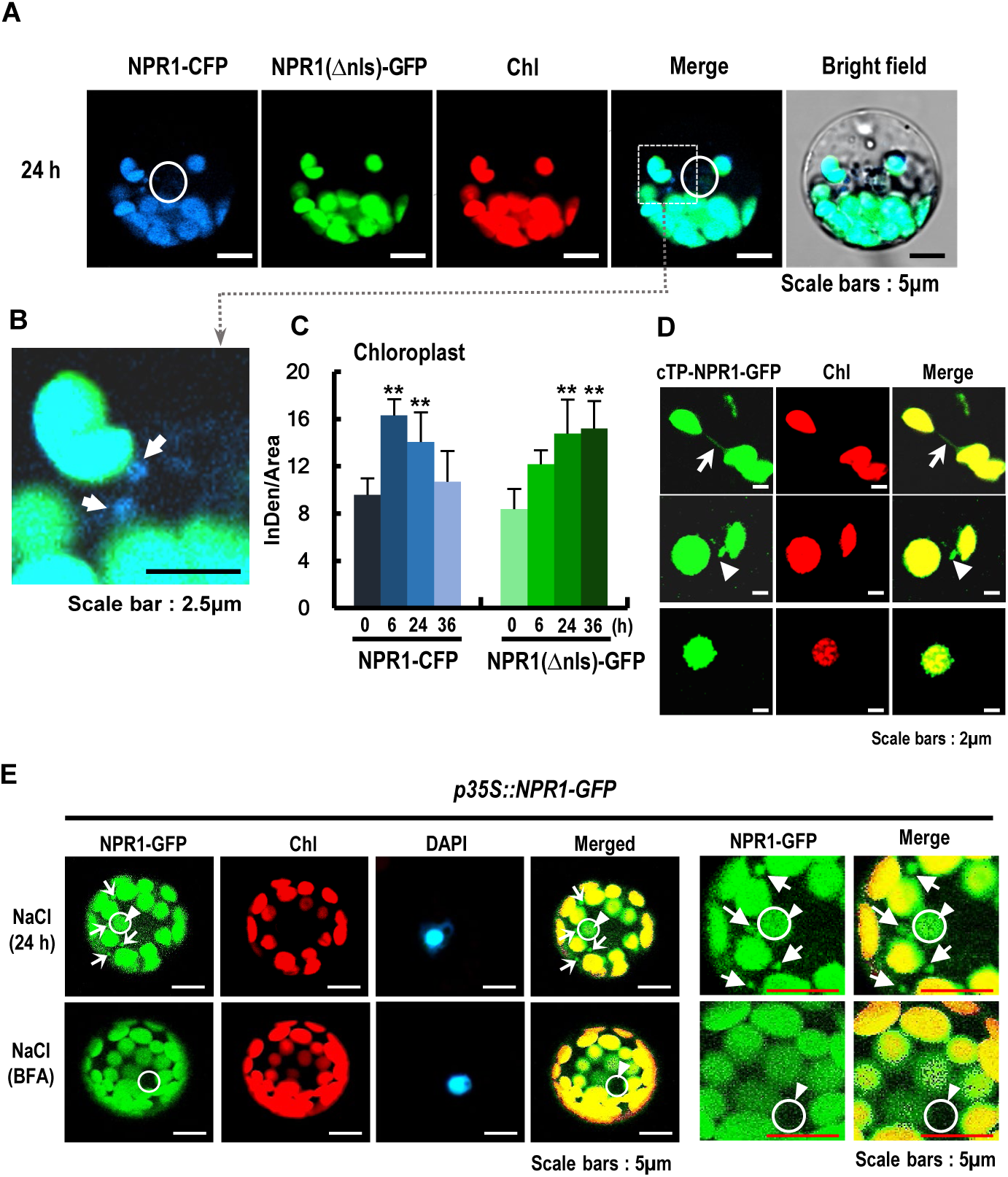
Signaling machinery from the chloroplasts to the nucleus under salt stress. **(A and B)** CLSM images observed after co-transient expression of *p35S*-driven *GFP*-tagged *NPR1* in which the nuclear localization sequence (NLS) was deleted and *p35S*-driven NPR1-CFP in mesophyll protoplasts of WT at 24 h after salt stress treatment. The white circle in the merge column indicates the nucleus site, and the blue NPR1-CFP is weakly visible in the nucleus (**A**). The dotted gray box in the merge column shows an image of the vesicle-shaped NPR1-CFP protruding from the chloroplast, which is enlarged (**B**). **(C)** Fluorescence intensity of NPR1-CFP (blue bar) and NPR1(△NLS)-GFP (green bar) in the chloroplasts after co-transient expression of both constructs. An asterisk indicates a significant difference between stress-treated or untreated cases (\*\**P* < 0.01). **(D)** CLSM images of stromules (upper row), cytoplasmic vesicles (middle row), and chloroplast protrusions (lower row) from isolated chloroplasts from *p35S::cTP-NPR1-GFP* transgenic plants under salt stress. White arrows indicate stromules and white triangles indicate chloroplast protrusions. **(E)** CLSM images of NPR1-GFP fluorescence in salt-stressed protoplasts treated with brefeldin A (BFA, bottom). Fifth and sixth column: enlarged CSLM images. White circles indicate the whole nucleus. White arrows indicate cytoplasmic vesicles.

Next, we investigated translocation vehicles that chloroplast NPR1 proteins translocate to the nucleus. In the experimental system under salt stress, rapidly moving vesicles emitting GFP fluorescence were observed in guard cells (Figure 1D and Supplemental Movie 3) and mesophyll cells (Supplemental Movie 2). In particular, rapid movement of vesicles containing NPR1-GFP molecules was observed around intact chloroplasts and in the perinuclear region of protoplasts from salt-stressed mesophyll cells (Figure 1B), suggesting intracellular trafficking of NPR1-GFP to the nucleus. Protrusions and vesicles from chloroplast bodies showed GFP fluorescence in isolated chloroplasts from leaves of salt-stressed *p35S::cTP-NPR1-GFP* tobacco transformants (Figure 4D).

These narrow, tiny structures were considered to be stromules, which are stroma-containing tubules that emanate from the main chloroplast body *in vivo* (Caplan, 2015). Stromules emanate from plastids at varying frequencies, which differ among environmental conditions and cell types (Ritzenthaler et al., 2002). Proteins, ROS, and other molecules flow through stromules, which might transport retrograde signaling molecules from chloroplasts to the nucleus (Caplan, 2015). Stromules may be a source of plastid-derived vesicles for signaling of environmental stimuli or for recycling of chloroplast contents (Ritzenthaler et al., 2002). We detected many fluorescent vesicles containing NPR1-GFP molecules in response to salt stress in leaves of tobacco overexpressing NPR1-GFP (Figure 1B and 1C). Although the precise role of stromule-derived vesicles or stromule-related chloroplast protrusions is unknown, they are suggested to play an important role in translocation of signaling components from chloroplasts to the nucleus.

Brefeldin A, which disrupts ER- and Golgi-mediated vesicular trafficking (Selga et al., 2010), reduced vesicle movement accompanied by significant reduction of NPR1-GFP fluorescence intensity in the nucleus under salt stress (Figure 4E). Therefore, stromule-derived vesicles may function as vehicles of NPR1 during chloroplast retrograde signaling by plastid–nuclear complexes through Golgi bodies, ER, and the nuclear envelope (Brigelius-Flohé and Flohé, 2011).

For ROS to affect gene expression, the oxidization of cysteine residues may modify the protein structure, resulting in larger oxidation products with disulfide bonds, or change enzymatic activity for production of signaling metabolites (van Eerden et al., 2017). The molecular weights of NPR1-GFP proteins of various sizes, including oligomeric forms >400 kDa, a tetrameric form, and a dimeric form were significantly and rapidly increased in chloroplast stroma proteins from leaves of salt-stressed *pNPR1::NPR1-GFP* and *p35S::cTP-NPR1-GFP* transformants (Figure 5A). In particular, the abundance of oligomers smaller than tetramers was significantly increased, suggesting that redox-sensitive NPR1 was converted to more reduced forms due to conformational changes in response to salt stress. Considering that cytosolic thioredoxins directly catalyze the NPR1 oligomer-to-monomer reaction (Tada et al., 2008), it is possible that stress-induced redox regulators markedly facilitated dimerization of NPR1 in chloroplasts, which was a more advantageous form to move. Only monomeric NPR1-GFP was detected in the nuclear fraction of *pNPR1::NPR1-GFP* transformants, which were maintained at significant levels at 6 to 48 h under salt stress (Figure 5B). Taken together, it is implied that translocation of chloroplast-localized NPR1 to the nucleus conveyed signals from the stressed chloroplasts and that NPR1 might be involved in retrograde chloroplast-to-nucleus signaling.

**Figure 5.**
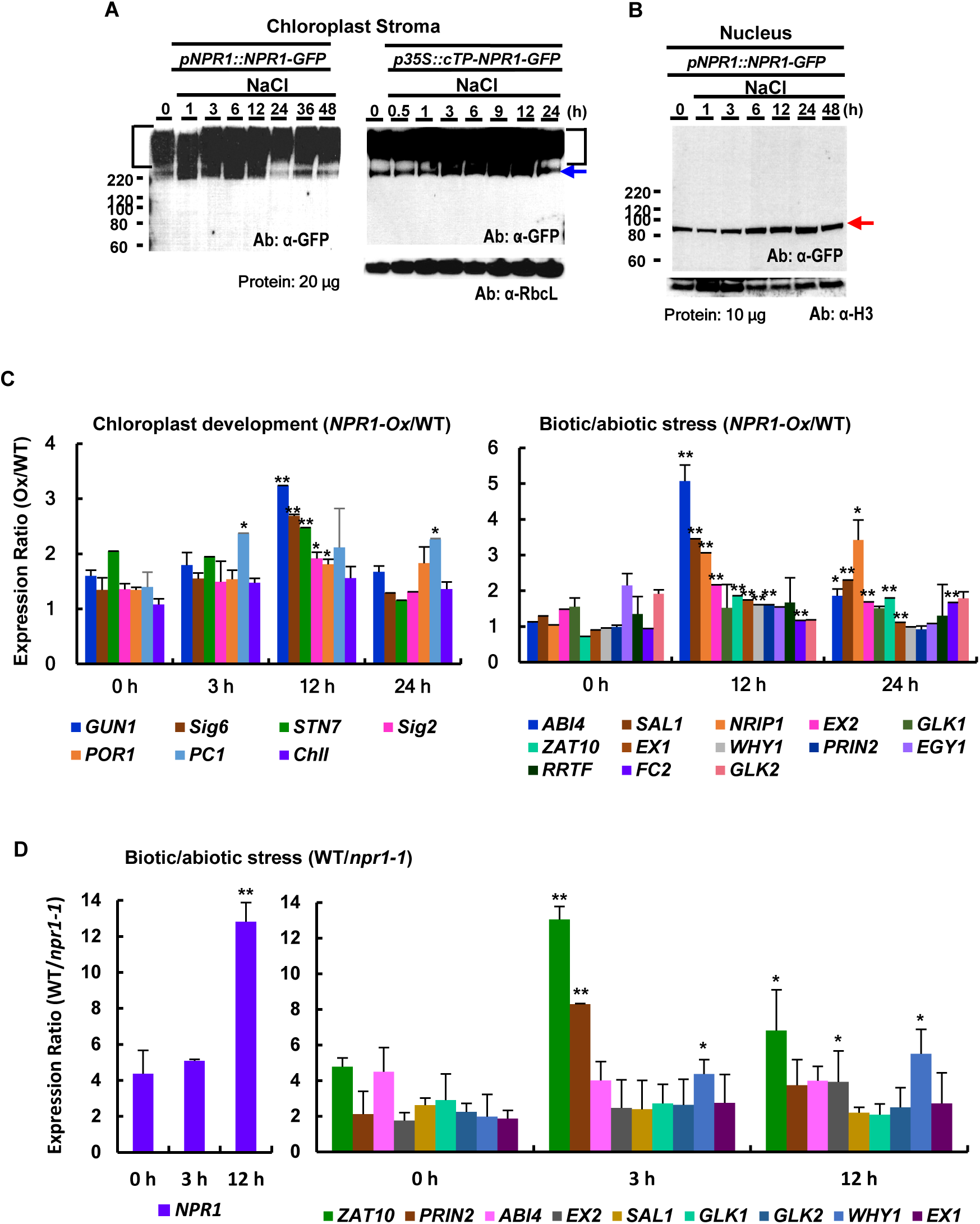
Western blot analysis of NPR1 for subcellular localization and expression analysis of retrograde signaling-related genes.

**(A and B)** Immunoblots showing NPR1-GFP in the protein fractions of chloroplast stroma (**A**) from *pNPR1::NPR1-GFP* (left) and *p35S::cTP-NPR1-GFP* (right) transgenic plants, and nucleus from *pNPR1::NPR1-GFP* (**B**) transgenic plants under salt stress by non-denatured SDS-PAGE. Oligomers (square bracket), dimeric form (blue arrow), and monomeric form (red arrow).

**(C)** Expression ratios of retrograde signaling-related genes for chloroplast development (left) and for biotic/abiotic stress (right) in *NPR1-Ox* versus wild type (WT) after salt stress.

**(D)** Expression ratios of retrograde signaling-related genes for biotic/abiotic stress in WT versus *npr1-1* mutant after salt stress treatment. The expression ratio was computed based on the relative expression level of each gene in *NPR1-Ox* versus WT

(**C**) or WT versus *npr1-1* mutant (**D**) after salt stress treatment. Chloroplast development: *GUN1*, Genomes uncoupled 1; *Sig6*, Chloroplast sigma factor 6; *STN7*, Serine/threonine-protein kinase 7; *Sig2*, Chloroplast sigma factor 2; *POR1*, NADPH:protochlorophyllide oxidoreductase; *PC1*, Plastocyanin; *ChlI*, Magnesium-protoporyphyrin chelatase subunit. Biotic/abiotic stress: *ABI4*, Abscisic acid-insensitive protein 4; *SAL1*, 3′-phosphoadenosine 5′-phosphate phosphatase; *NRIP1*, N receptor-interacting protein; *EX2*, Executer 2; *GLK1*, Golden 2-like 1; *ZAT10*, Zinc finger transcription factor 10; *EX1*, Executer 1; *WHY1*, single-stranded DNA-binding protein WHIRLY 1; *PRIN2*, Plastid redox insensitive 2; *EGY1*, ethylene-dependent gravitropism-deficient and yellow-green 1; *RRTF*, redox responsive transcription factor; *FC2*, ferrochelatase; *GLK2*, Golden 2-like 2; *NPR1*, Nonexpressor of pathogenesis-related genes 1. An asterisk indicates a significant difference between WT and *NPR1-Ox* tobacco or *npr1-1* Arabidopsis plants at an indicated time (\**P* < 0.05, \*\**P* < 0.01).

It is surprising that NPR1-GFP protein with a size of about 45 kDa (∼18 kDa in the C-terminal region of NPR1 and 27 kDa GFP), which has been designated CP45 (Seo et al., 2020), was detected in the nucleus (Supplemental Figure 5A and 5B). More importantly, nuclear CP45 was detected not only in NPR1-GFP transgenic plants, but also especially in plants expressing cTP-NPR1-GFP by non-reduced immunoblot analysis using a GFP antibody. The amounts of CP45 were significantly increased in the nuclear protein fraction after treatment with the proteasome inhibitors MG115 and MG132. These results indicated that the 45-kDa protein was regulated by proteasome-dependent degradation in the nucleus.

Using western blot analysis with an ubiquitin antibody after immunoprecipitation with a GFP antibody, we detected ubiquitinated NPR1 of the monomer and CP45 in the nuclear fraction of salt-stressed *p35S::cTP-NPR1-GFP* transgenic plants (Supplemental Figure 5C). These results confirmed that translocation of chloroplast-targeted NPR1 to the nucleus conveyed messages in the stressed chloroplasts and that NPR1 might be involved in retrograde chloroplast-to-nucleus signaling.

### Involvement of NPR1 in retrograde signaling communication

Although identification of retrograde signaling proteins remains elusive, several plastid proteins, including NB-LRR receptor-interacting protein 1 (NRIP1) and the single-stranded DNA-binding protein WHIRLY 1, are suggested to be involved in retrograde chloroplast signaling (Chan et al., 2016). Molecules involved in retrograde signaling communication were divided into two groups: components for chloroplast biogenesis and development, and components for stress response and immunity in plants. Under high salinity, the transcription ratio of each retrograde signaling-related protein for chloroplast development was above 1 in *NPR1* overexpression lines (*NPR1-Ox*) versus WT, in which the expression ratio was highest at 12 h as quantified using real-time RT-PCR (Figure 5C, left and Supplemental Figure 6A).

Stress-related retrograde signaling components, including transcription factors reported in the literature, were investigated next (Figure 5C, right). Transcript levels of these genes showed stress-inducible patterns in the WT and *NPR1-Ox*, increasing transiently and peaking at 12 h of salt stress (Supplemental Figure 6B), supporting the conclusion that the retrograde signaling-related responses were transient and short-lived. The transcription ratio of each retrograde signaling-related protein for the stress response was above 1 and the ratio increases were greatest at 12 h in *NPR1-Ox* versus WT (Figure 5C, left) implying that the transcription ratio profile is consistent with the localization pattern of nuclear NPR1 under salt stress.

To further explore the physiological functions of NPR1 in response to salt stress, we compared Arabidopsis WT and *npr1-1* mutant plants. Compared with the WT, the *npr1-1* mutant under salt stress showed reduced *F*_v_*/F*_m_ (Supplemental Figure 7A). The pattern for lower *F*_v_*/F*_m_ ratio was consistent with the gene expression characteristics of several chloroplastic components for photosynthesis (Supplemental Figure 7B). Salt stress significantly decreased the expression level of photosynthesis-related genes encoded in chloroplasts, but the decrease was more severe in the *npr1-1* mutant, implying that NPR1 functions in the positive regulation of gene expression in chloroplasts.

To investigate whether defective NPR1 affects expression of nuclear genes associated with retrograde communication, qRT-PCR analysis was performed. Under high salinity, expression levels of almost all analyzed genes remained similar to the basal level in the *npr1-1* mutant during the entire period of salt-stress treatment, which contrasted strongly with the gene expression pattern in WT plants (Supplemental Figure 8). The transcript level of all analyzed genes involved in stress-related retrograde signaling was significantly higher in the WT than in the *npr1-1* mutant. The expression ratio of each retrograde signaling-related gene was above 1 in WT versus *npr1-1* during the entire period of salt-stress treatment (Figure 5D). Therefore, these results indicated that NPR1 might be a major regulator in retrograde signaling pathways.

## Discussion

The chloroplast acts as a sensor of environmental and developmental cues that affect photosynthesis, relaying the information to the nucleus for coordination of plant growth and development and stress responses (Chan et al., 2016). This retrograde signaling regulates nuclear gene expression in response to developmental status and abiotic/biotic stresses (Fang et al., 2019). Recent advances have proposed a number of chloroplast retrograde signals, including carotenoid derivatives, isoprenoid precursors (methylerythritol cyclodiphosphate), 3′-phosphoadenosine 5′-phosphate, tetrapyrroles, heme, and ROS, together with transcription factors (Chan et al., 2016; Xiao et al., 2012; Pornsiriwong et al., 2017; Zhao et al., 2019a). These signals and related pathways build a communication network to regulate gene expression, miRNA biogenesis, RNA editing, and gene splicing to improve adaptation to developmental and stress stimuli (Fang et al., 2019; Petrillo et al., 2014; Godoy Herz et al., 2019; Zhao et al., 2019b; Zhao et al., 2020). Although several retrograde signaling modules have been identified, understanding the true complexity of the regulation of this pathway is in its infancy. Given that the regulation of retrograde signaling has been only partially explained, little is known about how signals are perceived and transmitted to the nucleus (Zhao et al., 2019b).

In this study, we showed that stress-induced chloroplast NPR1 was translocated to the nucleus in a redox-dependent manner via cytoplasmic vesicles or stromules (Figure 1, 2, and 4). In transformants of chloroplast-targeted NPR1-GFP fused with cTP, NPR1-GFP was detected in the nucleus under salt stress by immunoblotting and fluorescence image analysis, suggesting that NPR1 is moved from chloroplasts to the nucleus. Overexpression of chloroplast-targeted NPR1-GFP significantly enhanced stress tolerance and photosynthetic capability, and reduced accumulation of ROS under high salinity, compared with those of the WT (Figure 3). However, when NPR1 targeted to chloroplasts was overexpressed, ROS accumulation in the whole plant decreased but nuclear ROS accumulation increased (Figure 3D). This is considered to be because chloroplast-to-nucleus retrograde signaling was increased in accordance with ROS signaling.

The movement of chloroplast-targeted NPR1 to the nucleus was significantly enhanced by treatment with H_2_O_2_ and ACC (Figure 2C). It was previously reported that stress-induced NPR1 localization in chloroplasts was significantly reduced in transgenic plants harboring a silenced ACC synthase gene (*NtACS4* and *NtACS1*) and in transgenic antisense plants expressing NADPH oxidase genes (*RbohD* and *RbohF*) (Seo et al., 2020). Taken together, it is suggested that NPR1 translocation into chloroplasts is triggered by production of stress-induced ROS and ethylene, after which chloroplast NPR1 moved to the nucleus to act as a transcription coactivator (Spoel et al., 2009). This retrograde signaling induced the resultant transcriptional changes to contribute to attenuation of photosynthetic capability loss, alleviation of cell damage, and enhanced stress tolerance.

Given that NPR1 is a redox-dependent protein, it has been proposed that it may be involved in retrograde signaling pathways (Gläßer et al., 2014; Kleine and Leister, 2016). In particular, it has been suggested that the redox state of the photosynthetic electron transport chain triggers the movement of WHIRLY1 from the chloroplasts to the nucleus, and draws a parallel with the regulation of NPR1 from the cytosol to the nucleus (Foyer et al., 2014). In addition, β-cyclocitral or WHIRLY1, which are retrograde signaling components, have been suggested to increase SA synthesis and, as a result, NPR1 is connected to retrograde signaling through the translocation from the cytoplasm to the nucleus (Maruta et al., 2012; Lin et al., 2020). In the present study, the expression of genes associated with well-known retrograde signaling components increased transiently in transgenic tobacco lines (*NPR1-Ox*) compared with that of the WT (Figure 5C), and the expression of these genes in the Arabidopsis *npr1-1* mutant remained almost at basal levels (Figure 5D). These results suggested that NPR1 was directly linked to retrograde signaling pathways rather than SA-mediated translocation of NPR1.

For example, WHIRLY1, which belongs to a small plant-specific family of DNA/RNA binding proteins, has been proposed to move from the chloroplast to the nucleus in response to environmental cues, such as high light intensity (Foyer et al., 2014; Świda-Barteczka et al., 2018). However, it did not seem to be translocated from chloroplasts to the nucleus, but rather it was more likely that the WHIRLY1 protein comprised two isoforms localized in the chloroplasts and the nucleus, respectively (Lin et al., 2019). The dual functions of WHIRLY1 may be associated with its dual localization for coordination of the retrograde signaling from plastids to the nucleus (Ren et al., 2017). The plastid WHIRLY1 isoform predominantly affects stress-related gene expression, whereas nuclear WHIRLY1 primarily controls developmental gene expression. A shift from nuclear to plastid isoforms promotes H_2_O_2_ accumulation and accelerates plant senescence and SA accumulation (Lin et al., 2019; 2020). However, the regulatory mechanism governing the functional switch of WHIRLY1 for mediation of plastid-to-nucleus retrograde signaling remains unknown.

Our previous findings suggested that NPR1 undergoes a functional switch from a molecular chaperone in chloroplasts for emergency restoration, which is associated with proteostasis and redox homeostasis, to a transcriptional coactivator in the nucleus for adaptation to stress (Seo et al., 2020). NPR1 and WHIRLY1 show similarities in that both proteins exhibit dual functions as well as dual localization in the chloroplasts and nucleus. However, there are major differences between these two proteins. NPR1 is first imported to the chloroplast and then moves to the nucleus, whereas WHIRLY1 is considered to exist as two isoforms (Lin et al., 2019).

There are several explanations for the expression of different protein isoforms with different functions, which are generated by alternative splicing or different modifications in their respective compartments under exposure to stress. The redox state of the PQ pool in chloroplasts initiates an unknown chloroplast-to-nucleus retrograde signal to regulate the alternative splicing of nuclear genes through Pol II elongation (Godoy Herz et al., 2019). It is possible that alternative splicing of a certain gene can lead to the production of a protein with chloroplast/nucleus dual localization and this protein may act as a signaling protein of retrograde signaling. The chloroplast PQ pool redox state is indicated to connect chloroplast retrograde signaling with alternative splicing of nuclear genes (Petrillo et al., 2014; Jung and Mockler, 2014). We observed that increase in the DCMU-induced oxidized PQ pool and decrease in ROS production in response to DPI treatment were responsible for almost complete inhibition of nuclear NPR1 (Figure 1E). These results suggested that the redox states with PQ pool and ROS accumulation also affect the retrograde signaling of NPR1 from chloroplasts to the nucleus. Taken together, factors such as the chloroplast redox state that affect retrograde signaling influenced the accumulation of NPR1 in the chloroplasts and nucleus, which indicates that NPR1 moves from the chloroplasts to the nucleus.

Norflurazone treatment significantly increased nuclear NPR1 (Figure 1E) and chloroplast NPR1 (Seo et al., 2020), which are indicative of the enhancement of retrograde signaling from chloroplasts to the nucleus. These results are consistent with the report that norflurazone affects plastid RNA edition, which triggers retrograde signaling through the GENOMES UNCOUPLED 1 (GUN1)-mediated pathway (Zhao et al., 2020). GUN1, an integrator of multiple retrograde signaling pathways, is associated with plastid protein homeostasis, chloroplast protein import/cytosolic folding stress, and plastid RNA editing under stress (Wu et al., 2019). Stress-enhanced chloroplast NPR1 also participates in protein homeostasis assuming the role of a chaperone under salt stress, which also can activate protein quality control in plastids (Seo et al., 2020). Lin treatment was completely inhibited by translocation of nuclear NPR1 from chloroplast-targeted NPR1 (Figure 2D), linking the chloroplast’s function and retrograde signaling for tight control of proper allocation under stress. Therefore, one possible explanation for localization of NPR1 protein is the additional connections between retrograde signaling and its translocation for functional switch. Retrograde signaling of NPR1 triggered by redox regulation or stress-induced components in organelles are important regulatory mechanisms for plants to cope with environmental stresses. Plastid-to-nucleus retrograde signaling crucially contributes to normal growth and development in plants. For adjustment of cellular metabolism under adverse environmental conditions, particularly in photosynthetically active leaf cells, chloroplast NPR1 may be an emergency device, after which it functions as a retrograde communicator for the protective machinery from chloroplasts to the nucleus in a redox-mediated manner.

## Methods

### Plant Materials and Growth Conditions

*Nicotiana tabacum* cv. Wisconsin-38 was used for wild type (WT) and transgenic plants. *Arabidopsis thaliana* Col-0 and mutant *npr1-1* (Arabidopsis Biological Resource Center, Ohio State University, USA) were used in this study. The surface-sterilized seeds of tobacco and Arabidopsis were cultured on solid Murashige and Skoog (MS) medium (pH 5.8) under light (16L/8D, 100 μmol photons m^−2^ s^−1^) at room temperature (25 ± 5°C). After antibiotic selection, fully matured WT and transgenic plants were subjected to either salt stress (200 mM NaCl) or other chemicals. Solutions with salt and other chemicals were applied to the whole leaves with petiole or stems with several leaves in 20 mM MES buffer under light (100 μM photons m^−1^s^−1^) at 25 °C. For the mock treatment, tobacco petioles or stems were treated with MES buffer without salt stress.

### Gene constructs and transgenic plants

The preparation of the *p35S::NPR1-GFP* and the *p35S::NPR1-Ox* transgenic plants has been described previously (Seo et al., 2020). The open reading frame (ORF) of *NPR1* was PCR-amplified, and the resulting product was cloned into *pMBP* vector harboring *35S* promoter-driven green fluorescence protein (GFP) gene and *NOS terminator*. The native *NPR1* promoter from the genomic DNA of *Nicotiana tabacum* was amplified by PCR. For the *pNPR1::NPR1-GFP* transgenic plants, the 0.8 kbp DNA fragment of the *NPR1* promoter was PCR-amplified and cloned into the *promoter-less NPR1-GFP* construct, which was prepared from *p35S::NPR1-GFP* after deletion of the 35S promoter fragment. For the *p35S::cTP-NPR1-GFP* transgenic plants, the 237 bp PCR product of the transit peptide (79 amino acid residues) from the small subunit of RubisCo (GenBank AY220079) was cloned between the end of the *35S* promoter and the 5’ end of *NPR1* in the *p35S::NPR1-GFP* construct. For the *p35S::NPR1(Δnls)-GFP* construct, 54 bp of nuclear localization sequence (NLS, nucleotide position from 1,612 to 1665) in the *NPR1* gene was deleted from *p35S::NPR1-GFP* using a GeneArt Site-directed Mutagenesis PLUS kit (Thermo Fisher Scientific, USA).

The resulting plasmid constructs were introduced into an *N. tabacum* by *Agrobacterium* (strain LBA 4404)-mediated transformation. Homozygous T3 plants were used for further study in all cases of *NPR1-Ox*, *pNPR1-NPR1-GFP*, *p35S::NPR1-GFP*, and *p35S::cTP-NPR1-GFP*. Even if they were T3 homozygous lines, they were used as experimental plants after confirming kanamycin resistance. The surface-sterilized transgenic seeds were cultured on solid Murashige and Skoog medium (pH 5.8) under light (16L/8D, 100 μmol photons m^−2^ s^−1^) at room temperature (25 ± 5°C).

### RNA Isolation and Real-Time qPCR

Total RNA isolation was performed using Trizol Reagent (Molecular Research Center, USA). To analyze relative transcription levels by real-time qPCR, 1 μg of the total RNA from the leaves was reverse-transcribed for 30 min at 42°C in a 20-μl reaction volume using a High Fidelity PrimeScriptTM RT-PCR kit (Takara, Japan) according to the manufacturer’s instructions. The gene-specific PCR primers for qPCR, whose sequence information was obtained from the GenBank database, were designed according to a stringent set of criteria (Supplemental Table 1), including a predicted melting temperature of 60°C ± 5°C, primer lengths of 20 to 24 nucleotides, guanine-cytosine content of 50 to 60%, and PCR amplicon lengths of 100 to 250 bp. Real-time qPCR was performed in optical 96-well plates using a TP950 (Takara, Japan). Fluorescence threshold data (Ct) were analyzed using Thermal Cycler Dice Real-Time System Software (Takara, Japan) and then exported to Microsoft Excel for further analysis. The relative expression levels in each cDNA sample were normalized to the reference gene β-actin. The transcription levels were expressed relative to the reference gene β-actin after qPCR. The mean levels of relative mRNA expression for each gene in WT, overexpressing transgenic plants (*NPR1-Ox*), and *npr1-1* mutants were obtained. The expression ratio for each gene was calculated in WT versus *NPR1-Ox* plants or WT versus *npr1-1* mutants. The profile of the transcription levels was measured in genes, which involved retrograde communication for chloroplast development and operational signaling to abiotic/biotic stress under salt stress.

### Trypan Blue Staining

To monitor plant cell death, salt-treated tobacco leaf discs were immersed for 1 min in a boiling solution consisting of 10 ml of lactic acid, 10 ml of glycerol, 10 g of phenol, and 0.4% (w/v) trypan blue. After the plants had cooled to room temperature for 1 h, the solution was replaced with 70% (w/v) chloral hydrate. The stained plants were decolorized overnight and then photographed using a digital camera.

### Analysis of photosynthetic activity

Eight-week-old whole plants were transferred to a growth chamber, and steady-state net photosynthesis was determined on a Gas Exchange Measuring Station (Walz, Germany) using a built-in light source (210 *μ*mol photons m^−2^ s^−1^) (Wi and Park, 2002). A gas stream (60 l h^−1^, 21% O_2_, and 430 μl ^−1^ CO_2_) was provided continuously into the photosynthesis unit using a mass-flow control system. The leaf temperature was maintained at 25°C, and the humidity of the chamber was maintained at 70 ± 1%.

### Detection of GFP and CFP

The expression of NPR1-GFP or NPR1-CFP in the intact leaves and roots of stable transgenic *NPR1-GFP* transgenic plants or protoplasts prepared from transgenic *NPR1-GFP* tobacco plants was detected. The protoplasts were prepared by incubation in an enzyme solution (0.5 M mannitol, 1 mM CaCl_2_, 20 mM MES, 0.1% BSA, 1% cellulase R-10, and 0.25% marcerozyme R-10). The GFP fluorescence in the cells was detected using a confocal laser scanning microscope (FluoView 300, OLYMPUS, Japan and STELLARIS 8, Leica, Germany) or a fluorescence microscope (THUNDER Imager, Leica, Germany) equipped with a high-resolution CCD camera (OLYMPUS, FV300, Japan). GFP and CFP expression was visualized by excitation at 488 nm and emission at 520 nm and excitation at 450 nm and emission at 470 nm, respectively. Red chlorophyll fluorescence was visualized by excitation at 458 nm and emission at 647-720 nm. The fluorescence of DAPI (4’,6-diamidino-2-phenylindole) staining for the nuclei was visualized by excitation at 358 nm and emission at 461 nm. The fluorescence density was quantified using *ImageJ bundle software* (National Institutes of Health, USA).

### ROS Detection in leaves

For total ROS determination, leaf epidermal strips were peeled from tobacco leaves and floated on a solution of 50 μM 2’-7’dichlorofluorescein diacetate (DCFH-DA; Sigma Chemicals, St Louis, MO, USA). The leaf stripe samples were collected after salt stress treatment for the indicated time. The ROS was observed by fluorescence microscopy (excitation: 450 ± 490 nm; barrier 520 ± 560 nm) equipped with a cooled CCD camera (OLYMPUS, FV300, Japan). The superoxide anion level was determined using a nitroblue tetrazolium (NBT) solution (0.2%) in 50 mM sodium phosphate buffer (pH 7.5), and the H_2_O_2_ level was determined using diaminobenzidine (DAB) staining solution (1 mg/ml) in distilled water.

### Chloroplast and nucleus isolation, protein extraction, and Western blotting

To extract the total protein from tobacco roots, frozen tissues were ground to a powder and suspended in protein extraction buffer (50 mM Tris-HCl, pH 7.5, 150 mM NaCl, 5 mM EDTA, 0.1% Triton X-100, 0.2% Nonidet P-40 (NP-40), 50 μg/ml of tosyl-L-phenylalaninyl-chloromethylketone, 50 μg/ml of tosyl-L-lysine-chloromethylketone, serine protease inhibitors, 0.6 mM phenylmethylsulfonyl fluoride (PMSF), 80 μM MG115, 80 μM MG132, and one complete protease inhibitor cocktail tablet (Roche, USA)).

To extract the chloroplast stroma protein from the tobacco leaves, the chloroplasts were first isolated from the intact leaves using a chloroplast isolation kit (Sigma-Aldrich, USA), after which further intact chloroplasts were harvested using a 40/80% Percoll gradient. Intact chloroplasts were suspended in chloroplast lysis buffer (0.5 mM HEPES-KOH, pH 7.5, 2 mM MgCl_2_, 1 mM NaF, 1mM EDTA, 1 mM PMSF, 80 μM MG115, 80 μM MG132, and one complete protease inhibitor cocktail tablet (Roche, USA)). After lysate centrifugation, the supernatants were recovered as the total proteins or chloroplast stroma proteins. The inhibition of proteasome-dependent degradation was accomplished by 40 μM MG115.

To extract the nuclear proteins from the tobacco leaves, the nuclei were first isolated from the intact leaves, after which the nuclear proteins were extracted using a plant nuclei isolation/extraction kit, CelLytic^TM^ PN (Sigma-Aldrich, USA). The nuclei were collected from leaves using a nuclei isolation buffer by mesh filtering according to the manufacturer’s protocol. The cell lysate was prepared with 2.3 M sucrose by centrifugation at 12,000 x g for 10 min, after which the supernatant was removed. The nuclei pellet was then added to the nuclear protein extraction buffer. Nuclei proteins were then added to a working extraction buffer in addition to 80 μM MG115 and 80 μM MG132 and then centrifuged for 10 min at 12,000 x g. The pure supernatant was used to obtain soluble nuclear proteins.

Proteins (100 μg of the total proteins, 20 μg of chloroplast stroma proteins, or 20 μg or 50 μg of nuclear proteins) were separated by 4-12% Bis-Tris Plus (Novex, USA). The proteins were transferred onto iBlot 2 NC Regular Stacks (Novex, Israel), after which the blots were blocked using iBind Cards (Novex, Israel) according to the manufacturer’s instructions. NPR1-GFP proteins were detected by reacting the blots with the mouse monoclonal anti-GFP monoclonal antibody (Clontech, USA) and horseradish peroxidase-conjugated secondary antibody (Santa Cruz, USA). The bands were visualized using SuperSingnal West Substrate Working Solution (Thermo Scientific, USA) on X-ray film. The primary antibodies for anti-RbcL (Agrisera) and anti-histon3 (Agrisera) were used to confirm the equal loading of proteins.

### Immunoprecipitation using the anti-GFP antibody

For immunoprecipitation of the GFP-fused NPR1 proteins, the cytosol and nuclear proteins were extracted separately from the *p35S::cTP-NPR1-GFP* transgenic tobacco leaves using an immunoprecipitation buffer (1X phosphate-buffered saline, pH 7.4 (Cat. no. 10010-031, ThermoFisher Scientific, USA) containing MG115, MG132, and plant protease inhibitor cocktail (Sigma, USA). The protein lysates (30 µg) were precleared with 50 µl of sheep anti-rabbit magnetic beads in a microcentrifuge tube at room temperature for 1 h with gentle rotation. To the precleared lysate, 5% NGS (Normal Goat Serum, pH 7.4) in PBS was added for blocking. Subsequently, the primary anti-GFP antibody diluted in PBS was added to a final concentration of 0.2 µg/ml. After incubating the mixture at 4°C overnight with gentle rotation, the supernatant was discarded. The bead mixture was washed in wash buffer (5% NGS in PBS, 1% Triton® X-100, 3% BSA) by pipetting gently up and down. The bound proteins were eluted by boiling in 25 µl of 1X SDS sample buffer. The supernatant was analyzed by SDS-PAGE and immunoblotting with the anti-Ubiquitin antibody (Santa Cruz, USA).

### Transient expression in tobacco protoplasts

Mesophyll protoplasts from the WT or transgenic tobacco leaves were isolated using a protoplast extract enzyme solution (pH 5.7) consisting of 1% cellulose R-10 and 0.25% marcerozyme R-10. The leaf slices were transferred to a Petri dish containing enzyme solution and incubated in the dark for 12 h at 25°C. After incubation, the enzyme solution was discarded by mesh (10 mm) filtration, and the cells were overlaid with 1 ml of a W5 buffer (154 mM NaCl; 5 mM KCl; 125 mM CaCl_2_; 5 mM glucose; 1.5 M MES, pH 5.7). After gentle centrifugation (5 min at 80 g), the protoplasts floating at the interface were collected, washed with W5 (3/1 v/v), pelleted by centrifugation (10 min at 80 g), and resuspended in W5 solution. After stabilizing the protoplasts in ice for 30 min, protoplasts at a density of 10^6^/ml were used for a further transient transformation.

Protoplasts (300 µl) in W5 buffer were pipetted gently into a disposable 0.4 cm pre-chilled electroporation cuvette, and 50 µg of the DNA constructs in 10 µl of TE buffer was added. Electroporation was performed using the Gene Pulser Xcell System (Bio Rad, USA). Electroporation was carried out with 160 V/ 960 μF (voltage/capacitance) according to the manufacturer’s instructions. After electroporation, the cuvette was chilled on ice for 10 min, after which protoplasts were transferred to a conical tube using a glass Pasteur pipette with the addition of 500 μl of K3 media (154 mM NaCl; 125 mM CaCl_2_; 5 mM sucrose; 5 mM xylose; 1.5 mM MES, pH 5.7). These protoplasts were investigated using a confocal microscope.

### Statistical analyses

All experiments were repeated at least three times with three replicates, and the data from one representative experiment are presented. The statistically significant differences according to a t-test between the transgenic lines and respective controls at each time point are indicated with a single asterisk (*) (P < 0.05) or two asterisks (**) (P < 0.01).

## Acknowledgments

We thank Dr Choong-Min Ryu for providing seeds of the *npr1-1* mutant, and for critical advice and discussion of the experiments. We also thank Dr June M. Kwak for very helpful advice on this study. We thank Robert McKenzie, PhD, from Edanz Group (https://en-author-services.edanz.com/ac), for editing a draft of this manuscript. Funding: This work was supported by grants from the National Research Foundation of Korea (NRF2017R1D1A3B03034134 and NRF2020R1AC1012652) and the Korean Research Institute of Biology and Biotechnology (2019-0240 and 2020-0266) to K.Y.P. Competing interests: The authors declare no competing interests. Data and materials availability: All data are available in the manuscript or the Supplemental materials. The sequence of tobacco NPR1 was deposited in the GenBank database under accession number KY402167.

## Author contributions

S.Y.S. conducted all experiments. K.Y.P. designed and supervised the work, analyzed the data, and prepared the manuscript. Both authors discussed the results and approved the manuscript.

## Supplemental Information

Supplemental Table 1

Supplemental Figure 1-8

Supplemental Movie 1-3

